# Biomarker Quantification in Breast Cancer using Xenium In Situ

**DOI:** 10.64898/2025.12.08.692193

**Authors:** Amanda Janesick, Stephanie N. Kravitz, Weston Stauffer, Miriam Valencia, Sarah E.B. Taylor

## Abstract

Advances in spatial transcriptomics enable high-throughput quantitation of both established and novel biomarkers at single cell resolution, offering the potential to transform diagnostics. Using Xenium in situ technology in FFPE human breast samples, we address two challenges in the cancer field: 1. achieving reliable normalization of gene expression across heterogeneous sample populations; 2. identifying biomarkers that predict invasion or metastasis. We describe a scalable approach to identify low-variation housekeeping (HK) genes within any given sample set, then use those HK genes for cross- and intra-sample normalization of biomarkers. Analyzing 12 FFPE human breast samples–primarily ductal carcinoma in situ (DCIS)–with a custom 280-gene panel, we identified four HK genes (*EEF1G, EEF2, MALAT1*, and *RPLP0*) that exhibited minimal variability in tumor cells, four tumor cell biomarkers (*LDHA, SDC1, PIGR, SFRP1*) that increased or decreased with tumor grade, and one tumor-associated myoepithelial biomarker (*LAMC2*). Normalizing biomarkers to the four HK genes preserved the dynamic range of expression necessary for distinguishing tumor grades, outperforming HKs from legacy RT-PCR diagnostic panels. Lastly, we employed a cell-agnostic approach in the tumor periphery to quantify *MMP11*, a biomarker correlated with proliferative and potentially pre-invasive ducts. Our results establish a single cell normalization method for spatial in situ transcriptomics and reveal and quantitate biomarkers relevant to DCIS risk and progression.

## Background

Advancements in spatial transcriptomics and sequencing technologies are deepening insights into the molecular complexity underlying human diseases. The Xenium In Situ platform (10x Genomics) has been used in a variety of clinical research settings ^1^, providing highly sensitive, quantitative single cell gene expression data with accurate cell segmentation across a broad dynamic range ^2^. Ductal carcinoma in situ (DCIS) is a pre-invasive condition in human mammary tissue. Precise measuring of biomarker levels within the tumor cells, myoepithelial cells, or surrounding peritumoral stroma is essential for differentiating between indolent and aggressive forms of DCIS ^3^. Xenium has already shown strong performance in breast cancer FFPE samples, effectively distinguishing DCIS and tumor regions at different stages of cancer progression linked to distinct microenvironments ^4–6^. We extend this work to quantitatively assess biomarkers and correlate their expression with the cellular and genomic features of adjacent ducts.

A key challenge for Xenium and other similar technologies is the ability to directly compare, or normalize biomarker counts across different tissue samples. This ensures that changes in biomarker expression are due to the disease condition rather than variability in sample quality or preparation methods. One normalization approach is the inclusion of stable “housekeeping” (HK) genes, a strategy currently implemented in some clinical RT-qPCR-based cancer recurrence tests (e.g., Oncotype Dx). HK genes are typically invariantly expressed across all cells or cell types because they encode proteins essential for basic cellular functions and maintenance. The main limitation of the RT-qPCR tests, however, is that gene expression cannot be evaluated on a single cell-type basis. Bulk measurements mask variability between individual cell types, making it challenging to determine whether an HK gene genuinely exhibits stable expression across a heterogeneous sample. An advantage of using Xenium is its single-cell capability, which allows evaluation of genes that exhibit low variation within each major cell type. Here, we present a strategy for cross-sample normalization of Xenium data, followed by quantification and spatial profiling of biomarkers in cancer and myoepithelial cells and within the adjacent peritumoral stromal space.

## Results and Discussion

A robust normalization approach first requires an assessment of HK gene stability across a diverse array of samples. In this study, we analyzed 12 Formalin-Fixed Paraffin-Embedded (FFPE) human breast tissue samples: 8 ductal carcinoma in situ (DCIS), 3 invasive ductal carcinoma (IDC), and 1 normal (**Supplemental Table S1**). All samples were processed using Xenium with a fully custom gene panel that included breast tissue-relevant markers and 45 HK gene candidates for normalization approaches **(Supplemental Table S2**). Because tumor cells are inherently more transcriptionally active than other cell types (e.g., stromal cells), it is impractical to identify genes that maintain consistent expression levels across all cell types. Furthermore, certain applications may focus on a specific cell type, making it beneficial to identify stable genes with a defined cluster of cells.

Each breast cancer sample was independently clustered and major cell types were annotated. Gene stability within each section was assessed by calculating the coefficient of variation (CV) – standard deviation divided by the mean – across all cells within a given cell type group for each section. Some genes (*EEF1G, EEF2, MALAT1*) yielded low CV in all cell types we evaluated (tumor, myoepithelial, and T cells), while others were only stably expressed in specific cell types (**Figure 1A, S1A-B**). By contrast, differentially expressed genes exhibited high CV across tumor subtypes (**Figure 1B**). Similarly, low expressing genes experienced greater noise and stochasticity, making their inclusion in normalization strategies less feasible. This was true for some reference genes previously published in bulk RT-PCR assays such as *CCSER2, ANKRD17, PUM1, SYMPK, RER1*, and *TFRC* ^7–10^. Finally, we evaluated the five HK genes included on the Oncotype Dx gene panel and found that only two (*RPLP0* and *ACTB*) had low CV across the four cell types we evaluated (**Figure 1A, S1A-B**). *TFRC* and *GUSB*, in particular, exhibited significant variability across different tumor subtypes, indicating that their expression is influenced by tumor heterogeneity, making them unsuitable as HK genes.

**Figure 1.**
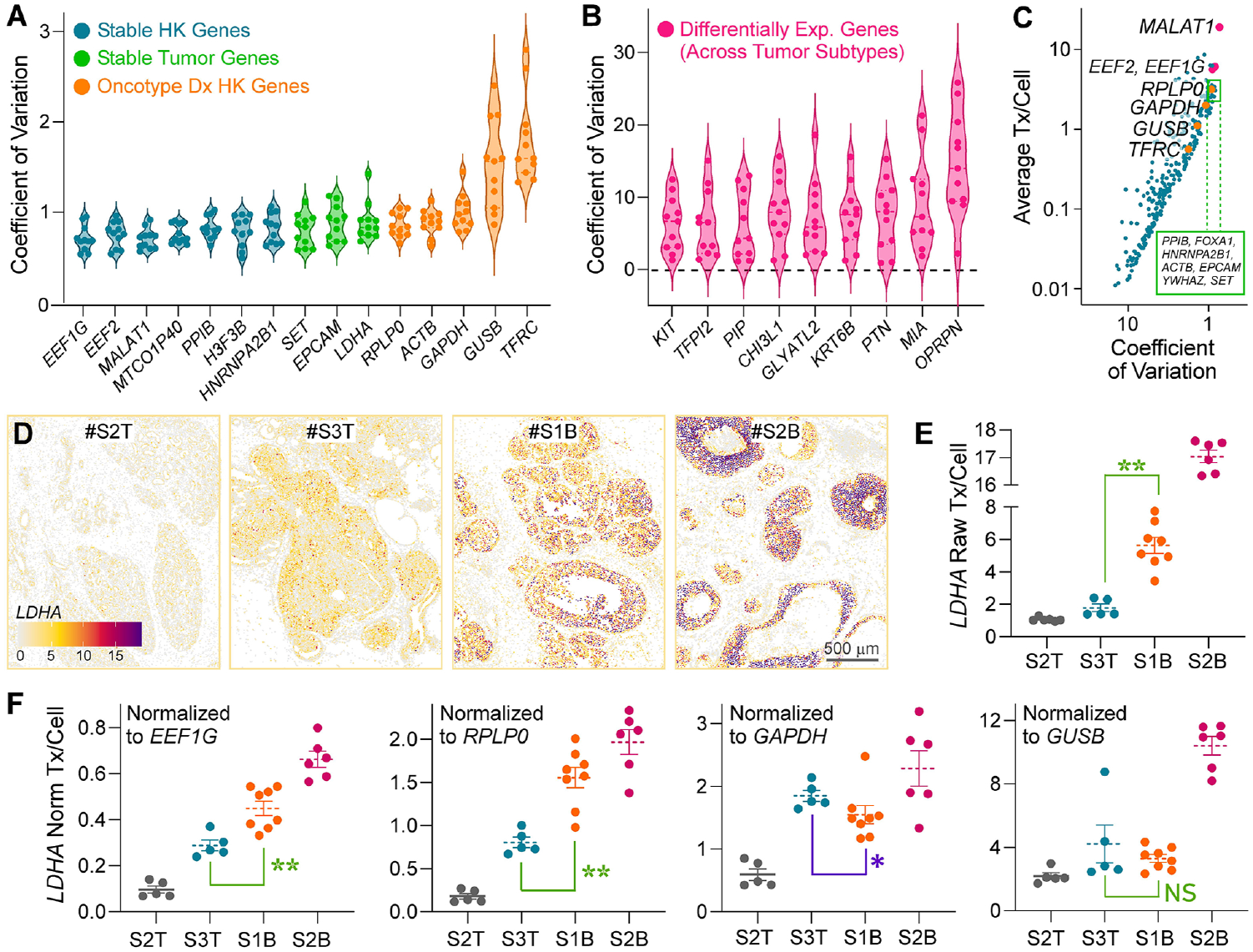
Evaluation of HK gene variation followed by *LDHA* normalization to selected HK genes across tumor grade. Eleven human breast cancer samples and one normal sample were independently clustered into major cell types. **(A, B)** Intra-section coefficient of variation (CV) among tumor cell clusters, where each dot within the violin plot is one section. Blue: Housekeeping (HK) genes with low CV for the cell type of interest. Green: Low CV genes not typically categorized as HK, and are generally cell type specific. Orange: Oncotype Ox HK genes. Magenta: Differentially expressed tumor genes yield high CV as expected. Example transcript localization plots are shown in **Supplemental Figure S2. (C)** Inter-section CV in tumor cells across all breast cancer tissue samples, plotted against average gene expression. Each dot denotes a gene on the Xenium panel. **(D)** Raw transcript counts for *LDHA* plotted as cell centroids in four different sections. **(E)** Average raw transcripts (tx) per cell and **(F)** normalized transcripts/cell for *LDHA*, where each dot is a tumor cluster. Dotted lines indicate the mean; error bars show S.E.M. The Mann-Whitney non-parametric test was used to assess statistical significance between #S3-Top and #S1-Bot tumor cell clusters.*= p ≤ 5 0.05.**= p ≤ 5 0.01. NS= not statistically significant.

We conducted comparisons of HK gene expression across all 11 breast cancer tissue samples (measuring inter-section heterogeneity) and found that *MALAT1, HNRNPA2B1, EEF1G, EEF2, SET, PPIB, GATA3, YWHAZ, EPCAM*, and *RPLP0* had the most robust expression and lowest CV in tumor cells (**Figure 1C**). We then performed the same inter-section calculation for myoepithelial cells and T cells. We provide a list of genes with stable CV (<1.5) across specific cell types, or across all three cell types we analyzed: tumor, myoepithelial, and T cells (**Supplemental Table S3**). We suggest researchers use this list to perform CV analyses on their specific sample sets to confirm reliable genes for their application. For the subsequent analyses in this study, we selected a subset of four genes (*EEF1G, EEF2, MALAT1*, and *RPLP0*) based on both our robust expression data and their strong support in existing literature, aiming for a universally applicable normalization set.

Next, we sought to use appropriate HK genes to retain meaningful differences in biomarker expression across all breast tissue sections. We classify potentially important biomarkers as those exhibiting a high Coefficient of Variation (CV) across tumor/epithelial cell clusters from the 12 sections (see **Figure 1B** for example). These genes are likely to effectively discriminate between distinct tumor cell populations within the breast cancer tissue. It is therefore essential to retain their full magnitude of expression differences when applying a normalization strategy. We calculated the CV for each gene using normalized expression values based on two different HK gene sets: 1. Oncotype Dx HK panel: *ACTB, GAPDH, GUSB, RPLP0*, and *TFRC*, or 2. Our selected HK genes: *EEF1G, EEF2, MALAT1*, and *RPLP0*. We found that using set #1 significantly decreased the overall CV compared to the raw count data (p < 0.0001) (**Supplemental Figure S3A**). In contrast, normalization using set #2 preserved the expression variation, yielding a CV that was comparable to the raw data (p = 0.0986) (**Supplemental Figure S3A**). These results demonstrate that conventional HK gene sets (Set #1) significantly suppress expression variance, potentially masking real biological results. In contrast, our refined panel successfully preserved gene expression effect size, validating its use for normalization.

One complication in pursuing HK normalization in tumor cells, where traditional biomarkers (*HER2, ER*, and *PR*) are expressed, is the Warburg effect ^11^. This phenomenon leads to a rise in tumor cell metabolic activity as tumor aggressiveness increases, thereby affecting some genes like *GAPDH* and *PGK1* which have historically been used as HK references ^12,13^. We demonstrate the importance of carefully selecting HK genes by investigating three DCIS and one normal sample. Among the DCIS samples, Section #S3-Top predominantly consisted of columnar cell hyperplasia (precursor to DCIS) and usual ductal hyperplasia (benign), and was therefore lower grade. Sections #S1-Bot and #S2-Bot were both comedo-type DCIS, with the latter being higher nuclear grade with necrosis. Although these HK genes still increase to some degree as the tumor becomes more aggressive, the magnitude is less pronounced compared to other HK genes that play a direct role in the glycolytic demand of tumor cells.

When examining raw transcript counts in tumor and epithelial cells, we observed that the expression levels of two genes, *LDHA* and *SDC1*, increased correspondingly with higher tumor grade (**Figure 1D-E, Supplemental Figure S3B**). *LDHA* facilitates the Warburg effect by promoting glycolysis, thereby supporting the metabolic needs and growth of cancer cells ^14,15^. *SDC1* has been associated with more aggressive tumor behavior and poorer prognosis ^16,17^. Two other genes, *SFRP1* and *PIGR*, decreased correspondingly with higher tumor grades (**Supplemental Figure S3C**). *SFRP1* inhibits the Wnt signaling pathway, while *PIGR* enhances immune surveillance; consequently, both genes are considered tumor suppressors. Since these genes serve as potential biomarkers for tumor grade, it is crucial to preserve their dynamic range and effect size during HK gene normalization.

We evaluated *LDHA* expression across the aforementioned four sections in tumor clusters normalized to low CV HK genes (*EEF1G, RPLP0*) versus higher CV HK genes (*GAPDH, GUSB*). Normalization to most HK genes maintained *LDHA* differential expression in low/mid-grade DCIS compared to high grade tumor (**Figure 1F**). However, some HK genes like *GAPDH* and *GUSB* failed to distinguish between a more benign tumor section (#S3-Top), and bona fide DCIS (#S1-Bot) (**Figure 1F**). Indeed, *GAPDH* caused a slightly reverse trend in *LDHA* expression. GAPDH is directly linked to the Warburg effect because it is the rate-controlling enzyme supporting conversion of glucose to pyruvate required by cancer cells ^18^. While *GUSB* is not directly linked to the Warburg effect, it is a lysosomal enzyme that can be induced as an adaptation to increased tumor malignancy ^19^. We argue that it is crucial to distinguish these more moderate cases of DCIS since they present the best opportunity for early intervention.

The progression of DCIS to invasive cancer depends on the breakdown of the barrier between ductal epithelial cells and the surrounding stroma ^20^. This barrier is structurally maintained by basement membrane (BM) components surrounding the duct, such as matrix metalloproteinases, collagen and elastin crosslinkers, fibulins, and heparan sulfate proteoglycans ^21^. We included a variety of BM genes on the Xenium panel and provided their cell localization in **Supplemental Table S4**. At the RNA level, these genes can be expressed by both luminal and basal myoepithelial cells, as well as the peritumoral stroma (i.e., fibroblasts, endothelial cells, and pericytes). These BM markers represent potentially important biomarkers because they can predict DCIS invasion prior to histological changes ^3^.

One key indicator of BM integrity in breast cancer is the loss of myoepithelial cells, which is quantifiable through standard histology. A discontinuous myoepithelial cell layer often serves as a proxy for increased tumor grade and DCIS risk ^22^. To this end, with a handful of markers, Xenium readily segments and identifies myoepithelial cells which can be evaluated across a tumor section ^4^. Recently, however, it has been acknowledged that directly measuring BM markers may yield more informative and predictive results when assessing DCIS risk ^23,24^. We evaluated myoepithelial cells for the expression of BM markers such as *LAMC2* and *COL17A1* in a DCIS tissue section (#S1-Top) containing normal-adjacent regions. We found that the BM gene *LAMC2* is significantly upregulated in tumor-versus normal-adjacent myoepithelial cells compared to other myoepithelial markers (*TAGLN, MYLK, COL17A1, KRT14*) (**Figure 2A-C**). This observation is consistent with studies showing decreased survival in breast cancer associated with *LAMC2* ^25^. A key mechanism involves the proteolytic cleavage of LAMC2 in a tumorigenic environment, producing fragments that weaken cell-matrix interactions and drive BM degradation ^26^. Consequently, our results suggest that *LAMC2* expression can serve as an early indicator of DCIS invasive potential, potentially detectable before the loss of myoepithelial cells becomes histologically apparent.

**Figure 2.**
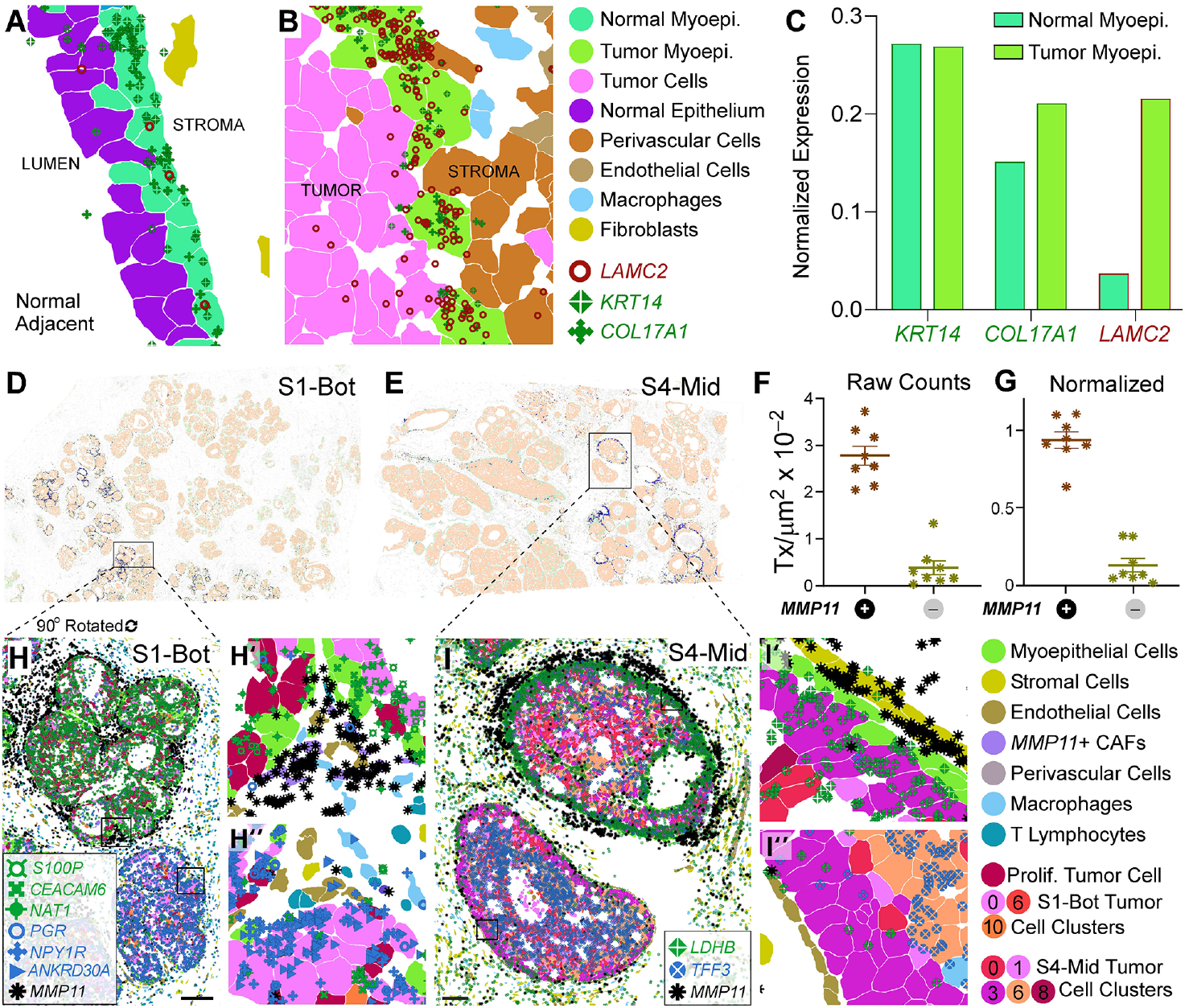
Exploration of biomarker quantification and normalization in the peritumoral space. **(A-C)** Section #S1-Top includes a region of histologically normal tissue adjacent to DCIS, enabling a comparison between non-cancerous ducts and those affected by tumorigenesis. **(A)** Normal epithelium (purple) is surrounded by myoepithelial cells (green) that express known markers such as *KRT14* and *COL14A1*, with minimal *LAMC2* transcripts detected. **(B)** In contrast, DCIS (pink) contains myoepithelial cells (green) that have upregulated *LAMC2* expression. **(C)** Quantitation of *KRT14, COL14A1* and *LAMC2* transcripts per myoepithelial cell adjacent to either normal or tumor duct, normalized to the geometric mean of *EEF1G, EEF2, RPLP0*, and *MALAT1*. **(D, E)** Spatial localization plots display *MMP11* expression (dark blue) overlaid on tumor (light orange), myoepithelial (light green) and stromal/immune (gray) cells in two sections. **(F)** Quantification of *MMP11* transcripts in 16 ROls (8 each from #S1-Bot and #S4-Mid) presented as transcript counts (tx) per µm_**2**_within a 30 µm peripheral zone around the duct. See **Supplemental Figure SGA** and **Supplemental Table SS** for the coordinate locations of all ROls included in the analysis. **(G)** *MMP11* expression from (F) normalized to the geometric average of *EEF1G, EEF2, RPLP0*, and *MALAT1* (tx/μm^2^ within the 30 μm periphery). (**H**) Region from #S1-Bot highlighting two ducts/ROIs (*MMP11*+ and *MMP11*−). Scalebar = 100 µm. (**H, H′**) Differential gene expression analysis demonstrates that *S100P, CEACAM6*, and *NAT1* are highly expressed in tumor cells adjacent to elevated peripheral *MMP11*. (**H, H′′**) *PGR, NPY1R*, and *ANKRD30A* are enriched in tumor cells within ducts surrounded by low *MMP11* expression. (**I**) Region from #S4-Mid showing two ducts/ROIs (*MMP11*+ and *MMP11*−). Scalebar = 100 µm. (**I, I′**) Differential gene expression analysis reveals that *LDHB* is localized to the DCIS perimeter adjacent to high peripheral *MMP11*. Scalebar = 100 µm. (**I, I′′**) *TFF3* is found a few tumor cell layers away from the DCIS perimeter in regions surrounded by low *MMP11* expression.

*LAMC2* is a relatively straightforward biomarker to quantitate on a per-cell basis since it is associated with well-segmented myoepithelial and tumor cells. By contrast, segmentation is less defined in the periphery of the duct where fibroblasts and endothelial cells have variable, elongated, and thin structures which are deeply embedded in extracellular matrix (ECM) making their boundaries more obscure. Some important BM markers, such as *LAMA2/4, LAMB1, MMP2/11/14, FBLN1, TIMP1*, and *LOXL1*, are expressed in the stroma; however, due to the dense ECM we observed some transcripts were not assigned to cells (**Supplemental Figure S4**). This leads to an underestimation of these potentially important biomarkers of DCIS risk when using a per-cell quantitation approach. This limitation is common to all spatial analysis platforms and is attributed to the inherent structural density of the ECM and stromal cell morphology. Therefore, we decided to employ a cell-agnostic strategy for capturing BM markers in the tumor periphery at a specified distance from the ductal cell boundary (**Supplemental Figure S5**).

Within the peritumoral space of DCIS, *MMP11* exhibited spatial heterogeneity and specificity; it is highly localized in the tumor periphery compared to other markers expressed more ubiquitously in the stromal milieu (**Figure 2D, E**). MMP11 protein degrades extracellular matrix components, facilitating tumor invasion, and modulates the tumor microenvironment to support cancer cell survival and metastasis ^27^. Although *MMP11* is expressed by a subset of endothelial cells and fibroblasts, we quantitated *MMP11* transcripts within a 30 µm-thick border surrounding the duct (see Methods), defined by the union of tumor and myoepithelial cell boundaries (**Supplemental Figure S5**). We compared four ducts with high *MMP11* and four ducts with low *MMP11* expression in two different sections (#S1-Bot and #S4-Mid; **Supplemental Figure S6**). *MMP11* transcripts quantitated in the periphery of each selected duct were expressed as transcripts/µm^2^ (**Figure 2F**) or as normalized values (using the HK genes selected above) (**Figure 2G**).

We next conducted cell type proportion and differential gene expression analyses on these ducts. In section S1-Bot, we observed that elevated *MMP11* periphery expression was correlated with a greater proportion of proliferating cells–positive for *MKI67, RRM2, TOP2A, CCNB1*, and *TYMS*) (**Supplemental Figure S6**). This finding is consistent with reports showing that *MMP11* expression in the tumor microenvironment is associated with increased bioavailability of IGF-1, a potent mitogen, as well as recruitment of immune cells expressing pro-proliferative cytokines such as IL-6/8, IL-1β, TNF-α, TGF-β, and CSF1 ^28^. In addition, S1-Bot ducts circumscribed by high *MMP11* expression were enriched for *CEACAM6, NAT1*, and *S100P* transcripts (**Figure 2H, H′**). Notably, *CEACAM6* and *S100P* are associated with more aggressive and invasive forms of breast cancer ^29–32^.

By contrast, we found that *MMP11*-negative ducts were associated with higher levels of *PGR, NPY1R*, and *ANKRD30A* (**Figure 2H, H′′**), suggesting a retention of normal glandular function, hormone-responsiveness, and therefore, a less aggressive phenotype ^33–35^. In section #S4-Mid, elevated MMP11 periphery expression was linked to an increase in *LDHB* expression, which was localized to the luminal periphery of the duct (**Figure 2I, I′**). Such spatial distribution of *LDHB* in conjunction with a microenvironment rich in *MMP11* suggests that cells in the leading edge are metabolically primed for invasion ^36,37^. By contrast, *TFF3* is found a few tumor cell layers away from the DCIS perimeter in regions surrounded by low *MMP11* expression (**Figure 2I, I′′**). In summary, a robust method for quantifying genes such as *LAMC2* and *MMP11* provides a direct, mechanistic readout of BM penetration and is, therefore, a critical indicator of tumor invasiveness.

## Conclusion

Leveraging Xenium in situ transcriptomics, we have established a single-cell normalization approach that overcomes the limitations of bulk assays and conventional housekeeping genes. Our study reveals a refined set of housekeeping genes (*EEF1G, EEF2, MALAT1, RPLP0*) which do not scale with the energy demands of tumor cells but instead remain relatively constant. We show that these HK genes preserve the dynamic range of biomarker expression, using the example of *LDHA*, necessary to distinguish tumor grades. These four HK genes are strong candidates for direct inclusion in the Xenium panel, given their fundamental cellular function and supporting evidence from both the literature and this study. However, we highly recommend an initial screening of a broader set of HK genes (**Supplemental Table S3**) and validating them in the specific tissue of interest to confirm their utility. We introduce novel indicators of invasive potential based on the disruption of BM markers—a requisite step for tumor cells to infiltrate surrounding tissue. *LAMC2* is found predominantly in tumor-adjacent myoepithelial cells; *MMP11* is found in the peritumoral stroma where its presence correlates with proliferative and aggressive ductal signatures. As the field of spatial transcriptomics advances, questions will focus more on biomarker quantitation rather than detection alone. The methodology we describe will be essential for accelerating our understanding of differential gene expression in complex systems and for the robust development of next-generation diagnostics.

## Acknowledgements

We would like to thank the 10x Genomics Xenium development team as well as Robert West and Mike Angelo (Stanford University) for their valuable advice regarding basement membrane genes and myoepithelial cells in the context of DCIS risk. We are grateful to Qiang Gong (10x Genomics) for generating Supplemental Figure S4 and Ian Fiddes (10x Genomics) for the Xenium custom probe design. Thank you to Juan Pablo Romero and Robert Shelansky (10x Genomics) for early discussions on normalization strategies. We also appreciate the thoughtful feedback on the manuscript draft provided by Michelli Faria de Oliveira, Corey Nemec, Andrew Gottscho, Marisa Lim, Byron Hartman, and Roman Yelensky (10x Genomics).

## Methods

### Biomaterials

Human breast FFPE-preserved blocks were obtained from Avaden, BioIVT, and Discovery Life Sciences. The sample set comprised 11 blocks representing a range of breast cancer stages and grades, including both DCIS and invasive lesions, as well as one normal breast tissue sample (see **Supplemental Table S1**). Additional pathology annotations, beyond those provided by the biobanks, were generated by Agoko (https://agoko.be/HOME.html).

### Tissue Preparation and Xenium Assay

Tissues were prepared following 10x Genomics demonstrated protocols “Xenium In Situ for FFPE - Tissue Preparation Guide” ([CG000578](https://www.10xgenomics.com/support/in-situ-gene-expression/documentation/steps/tissue-prep-ffpe/xenium-in-situ-spatial-profiling-for-ffpe–––tissue-preparation-guide)) and “Xenium In Situ for FFPE Tissues – Deparaffinization & Decrosslinking” ([CG000580](https://www.10xgenomics.com/support/instruments/xenium-analyzer/xenium-in-situ-for-ffpe-tissues–––deparaffinization-decrosslinking)). Probe hybridization, washing, ligation, amplification, and cell segmentation staining were performed following the “Xenium In Situ Gene Expression with Cell Segmentation Staining User Guide” ([CG000749](https://www.10xgenomics.com/support/in-situ-gene-expression/documentation/steps/assay/xenium-in-situ-gene-expression-with-morphology-based-cell-segmentation-staining-user-guide)). Instrument operation followed the procedures in the “Xenium Analyzer User Guide” ([CG000584](https://www.10xgenomics.com/support/in-situ-gene-expression/documentation/instruments/xenium-analyzer/xenium-analyzer-user-guide)). Automated data processing on the Xenium instrument was run with software analysis version 4.0.

### Serial Section IHC and Post-Xenium H&E

H&E staining was conducted according to the 10x Genomics demonstrated protocol “Xenium In Situ Gene Expression - Post-Xenium Analyzer H&E Staining” ([CG000613](https://www.10xgenomics.com/support/in-situ-gene-expression/documentation/steps/assay/xenium-in-situ-gene-expression-h-e-staining)). Stained slides were imaged on an Olympus VS200 scanner. The digital images were then converted from VSI to OME.tif format in QuPath-0.5.1, following the 10x Genomics tutorial “Converting Post-Xenium Images for Xenium Explorer Compatibility”. H&E images and alignment files for all 12 sections are available under the Output and Supplemental Files tab at: https://www.10xgenomics.com/datasets/xenium-ffpe-human-breast-biomarkers

### Gene Panel

A fully custom standalone Xenium panel of 280 genes was designed using the cloud-based Xenium Panel Designer v3.4.2 from 10x Genomics (**Supplemental Table S2)**. The panel enables comprehensive cell type identification in human breast cancer with particular emphasis on basement membrane (BM) markers (see **Supplemental Table S4**) and myoepithelial genes such as *ACTA2, MYH11, KRT14/17*, and *CNN1*. A number of human breast tumor biomarkers were included such as *ERBB2, PGR, ESR1, GATA3*, and *MKI67*. The panel also includes 45 housekeeping (HK) genes that were derived from the literature ^7–9^, the Oncogene Dx reference panel ^10^, and internal Flex or Xenium breast datasets. These genes were selected based on their low dispersion feature observed in these datasets. Some of the HK genes had probeset numbers reduced due to predicted high expression.

### Cell annotation of Xenium data

Xenium data was imported into Seurat v5.0.1.9007, without transcript coordinate data, according to the vignette, “Analysis of Image-based Spatial Data in Seurat” ^38^. Cells were retained for downstream analysis only if they contained more than 10 total transcripts and detected over five unique genes. We normalized the data using SCTransform prior to clustering, differential gene expression analysis, and spatial visualizations (e.g., Supplemental Figure S2). Using the `FindClusters`function (resolution = 0.3, dimensionality determined via elbow plot ranking of principle components), the major cell type groups (e.g., tumor/epithelial, myoepithelial, immune, endothelial, stromal/fibroblasts) were manually annotated. Additional subclustering was performed in one instance: myoepithelial cells from the S1-Top section were subdivided into normal-adjacent and tumor-adjacent populations. Cell annotation files for all 12 sections are available under the Output and Supplemental Files tab at: https://www.10xgenomics.com/datasets/xenium-ffpe-human-breast-biomarkers

### Evaluation of Housekeeping (HK) Gene Stability

For each of the annotated cell type groups, a raw count/barcode matrix was exported (using the `FetchData`function in Seurat). The intra-section coefficient of variation (CV) was calculated by dividing the standard deviation by the mean of HK expression values across all cells of that type within a given tissue section. CV was plotted in GraphPad Prism v10.3.1 for each cell type of interest. Certain sections were excluded from these plots for the following reasons: 1. The normal breast cancer section was excluded from Figure 1A, since normal epithelium is transcriptionally distinct from tumor epithelium; 2. Three invasive sections did not have myoepithelial cells, and therefore were not plotted in Supplemental Figure S1A. 3. We detected no T cells in two samples and therefore these were not plotted in Supplemental Figure S1B. Differentially expressed genes in Figure 1B were determined using the FindMarkers function in Seurat. Inter-section CV (Figure 1C) was calculated across tumor cells from all 11 breast cancer sections (excluding normal - #S2-Top).

### Normalization of Biomarkers to HK Genes

For each cluster and for every gene, raw transcript counts were exported (using the `AggregateExpression`function) and then divided by the total number of cells belonging to that cluster. This yielded an average transcripts-per-cell (tx/cell) value for each biomarker or HK gene. The rationale to aggregate counts at a cluster level was chosen because scatter plot analysis confirmed that housekeeping and biomarker gene expression did not co-vary at the single-cell level. Raw and HK-normalized transcript values per cell were visualized for each cluster using GraphPad Prism v10.3.1.

Normalization for Figure 1F was performed by dividing biomarker tx/cell by HK tx/cell. Statistical comparisons of tx/cell values between S3T and S1B blocks were performed using the Mann-Whitney nonparametric test. Normalization for Figure 2C was performed by dividing the tx/cell value for each myoepithelial marker by the geometric mean tx/cell value of four low-variance housekeeping genes—*EEF1G, EEF2, MALAT1*, and *RPLP0*—calculated separately for the normal-adjacent and tumor-adjacent myoepithelial cell clusters. For Figure 2G, normalization followed the same principle, but was based on spatial area rather than cell segmentation. *MMP11* transcript count was divided by the geometric mean of *EEF1G, EEF2, MALAT1*, and *RPLP0* counts, with all values expressed per μm^2^ within the designated periductal region (see “Tumor Periphery Analysis”). For Supplemental Figure S3A, the top 100 differentially expressed tumor cell genes (CV > 1, maximum expression in any single tumor cluster > 2 transcripts/cell) were normalized by dividing the tx/cell value by the geometric mean tx/cell value of either 1. four low-variance HK genes—*EEF1G, EEF2, MALAT1*, and *RPLP0* or 2. Oncotype Dx HK genes— *ACTB, GAPDH, GUSB, RPLP0*, and *TFRC*, across all tumor cell clusters. Statistical comparisons of CV between groups were performed using a matched sample one-way ANOVA followed by Dunnett’s multiple comparisons test.

### Tumor Periphery Analysis

To quantify BM markers such as *MMP11* in areas where a substantial number of transcripts are detected outside cell segmentation boundaries (**Supplemental Figure S4**), we employed a cell-agnostic approach that captures all transcripts within the tumor periphery.

We defined 16 regions of interest (ROIs) based on the presence or absence of the BM marker *MMP11*: 8 *MMP11*-high and 8 *MMP11*-low, from two different sections (**Supplemental Figure S6A, C**). Using Xenium Explorer v4.1, we used the lasso tool to roughly outline broad areas around the ducts of interest and exported the corresponding cell IDs from the ROIs (cell coordinates or cell stats.csv) (**Supplemental Figure 5A**). Although this approach is partly manual, it effectively reduced the search area for subsequent computational refinement of the tumor periphery. We then focused on isolating myoepithelial and tumor cell types within these ROIs to precisely delineate the duct regions. Then we employed the `cell_boundaries.parquet`file to generate segmentation masks for the selected ductal cells (**Supplemental Figure 5B**).

Using the R package `sf`(Bivand 2023) to access and manipulate simple spatial features, we buffered the cell boundaries to unionize polygons which constitute the duct defined by tumor and myoepithelial cells (shown in blue in **Supplemental Figure 5B)**. Since cells are not perfectly packed, we minimized gaps between them by applying a buffer of 5-20 μm (depending on the local gap density) to join the cells more smoothly together. After establishing the ductal perimeter geometry, we expanded the border by 30 μm to generate the tumor periphery space (displayed in light blue in **Supplemental Figure 5C**). We chose the 30 μm distance, accounting first for the eosinophilic acellular matrix (10-20 μm thick) and then extending to the first cell layer of CAFs adjacent to the matrix (∼10 μm). This distance also allowed us to capture the majority of *MMP11* transcripts in *MMP11*+ ROIs. The periphery mask was then used to isolate all transcripts overlapping the region, independent of cell segmentation.

In some instances, expanding the duct border by 30 μm causes the periphery to overlap with a neighboring duct. To avoid counting transcripts from these nearby tumor and myoepithelial cells, we identified and subtracted the area occupied by those neighboring cells from the periphery mask. This correction allowed us to accurately measure the periphery area in square micrometers (μm^2^) using the `sf`package’s `st_area()`function (see **Supplemental Figure 5D**).

To analyze transcripts within the 30 μm tumor periphery, we first retrieved the x and y coordinates for each transcript from the `transcripts.parquet`file. To streamline data processing, we restricted loading transcripts that fell inside the bounding box of the originally lasso’d region in Xenium Explorer—defined by xmin, ymin, xmax, and ymax coordinates that encompass the periphery area. Next, we used the `sf`package to identify which transcripts overlapped the precise boundaries of the actual tumor periphery region defined above. We excluded transcripts assigned to cell_ids corresponding to tumor or myoepithelial cells. For each gene of interest, transcript density within the periphery region of a given ROI was calculated by dividing the number of transcript counts by the area of the periphery region.

The tumor periphery script is provided in Supplementary Materials and the ROI coordinates are found in **Supplemental Table S5**. The only files required are: `cell_stats.csv`of the lasso’d ROI exported from Xenium Explorer; 2. Cell type annotation of the duct, 3. `cell_boundaries.parquet`and 4. `transcripts.parquet`

## Data Availability

The Xenium data generated in this study have been deposited at: https://www.10xgenomics.com/datasets/xenium-ffpe-human-breast-biomarkers

