## Supplemental Figures for "Biomarker Quantification in Breast Cancer using Xenium In Situ"

### Supplemental Figure S1

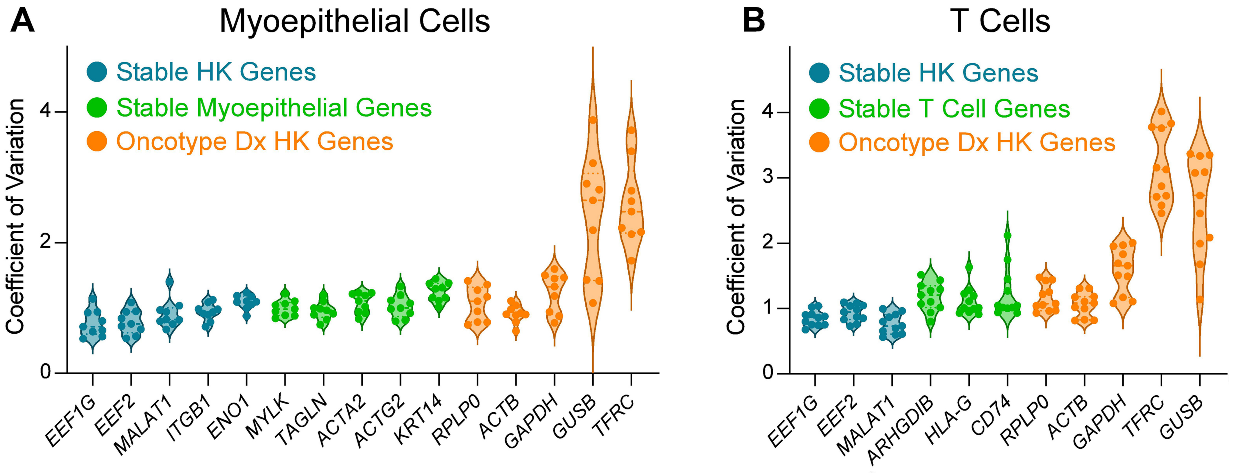


**Supplemental Figure S1. Evaluation of intra-section HK gene variation across myoepithelial and T Cell types in twelve human breast samples.** A total of 11 human breast cancer and 1 normal sample were independently clustered into major cell types. Myoepithelial and T cells were each represented by one cluster. For each major cell type (**A**: myoepithelial cells, **B**: T cells), the Y-axis displays the CV, where each dot within the violin plot represents one sample. Blue: HK genes with low CV for the cell type of interest. Green: Low CV genes not typically categorized as HK, and are generally cell type specific. Orange: Oncotype Dx HK genes. Example transcript localization plots are shown in **Supplemental Figure S2**.

### Supplemental Figure S2

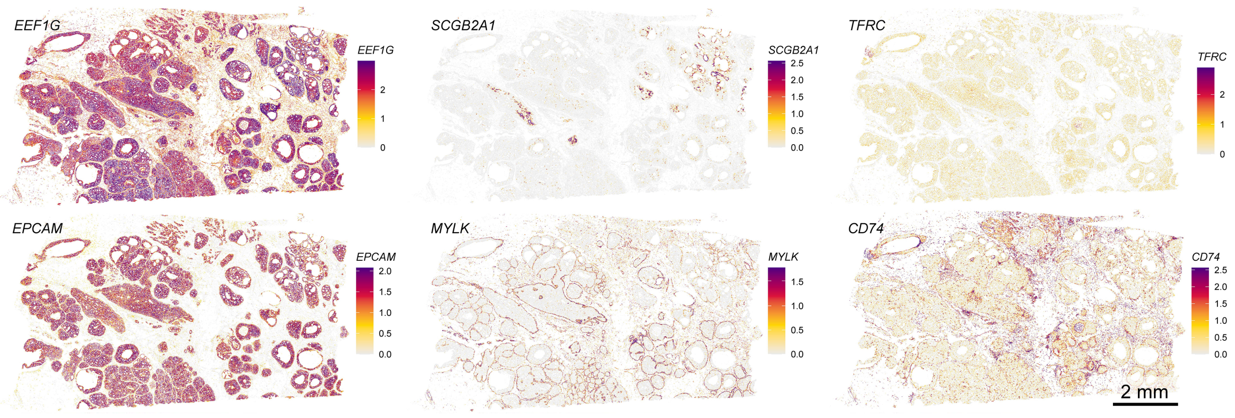


**Supplemental Figure S2. Example transcript localization plots of genes categorized in Figures 1A, 1B, S1A, and S1B.** Transcript counts (SCTransform normalized) are plotted as cell centroids in Section S4-Mid. *EEF1G* is an ideal HK gene (lowest CV). *SCGB2A1* is a differentially expressed gene (high CV). *TFRC* is an Oncotype Dx reference gene with low expression and high CV. *EPCAM*, *MYLK*, and *CD74* are robustly expressed, cell-type specific stable genes (low CV) for tumor, myoepithelial, and T cells respectively.

### Supplemental Figure S3

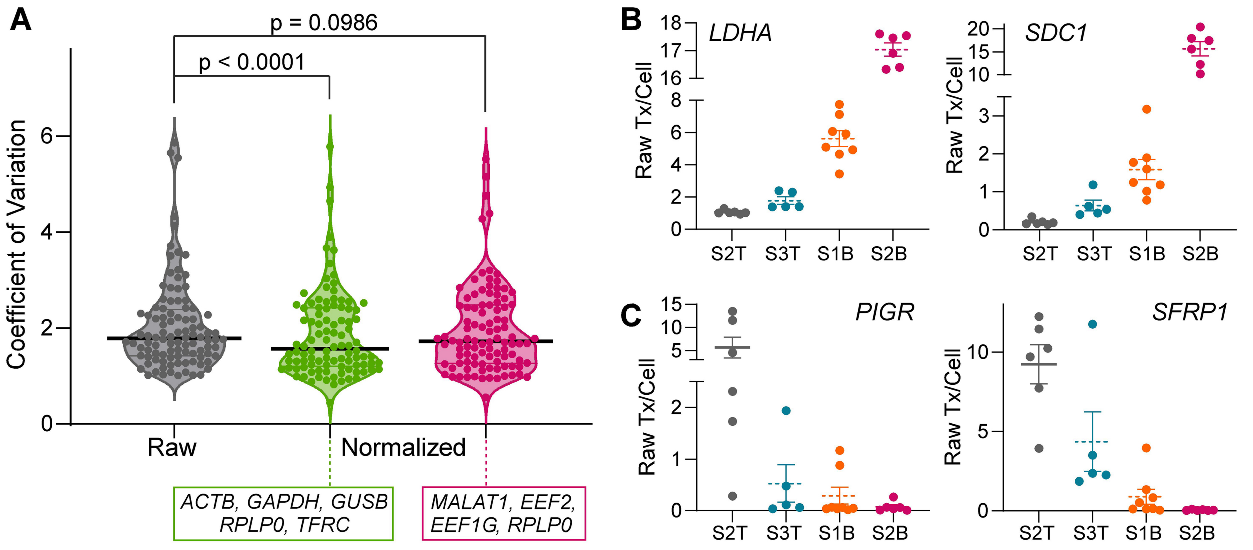


**Supplemental Figure S3. Effective normalization preserves biological heterogeneity important for tumor grade-correlated biomarkers.** Twelve human breast cancer samples were independently clustered into major cell types. **(A)** In 73 tumor/epithelial cell clusters across these 12 sections, we identified 100 differentially expressed genes (potential biomarkers). Raw transcripts/cell values for each tumor/epithelial cell cluster is shown in gray. We aggregated counts for each cluster and gene, then performed normalization with either set 1: *ACTB*, *GAPDH*, *GUSB*, *RPLP0*, and *TFRC* (green) or set 2: *EEF1G*, *EEF2*, *MALAT1*, and *RPLP0* (magenta). Thick black horizontal line is the mean and thin lines are the quartiles. The reported p-values are the results of a matched sample one-way ANOVA followed by Dunnett's multiple comparisons test. (**B, C**) Four human breast samples of increasing tumor grade: S2T is normal breast, S3T is columnar cell and usual ductal hyperplasia, S1B is DCIS, and S2B is DCIS with necrosis. In tumor/epithelial cells across these four sections, we observed two biomarkers with increased expression (*LDHA* and *SDC1*) and two biomarkers with decreased expression (*PIGR* and *SFRP1*). Average raw transcripts (tx) per cell for (**B**) *LDHA*, *SDC1*, (**C**) *PIGR*, and *SFRP1*, where each dot represents a tumor/epithelial cell cluster. Dotted lines represent the mean, and error bars indicate S.E.M.

### Supplemental Figure S4

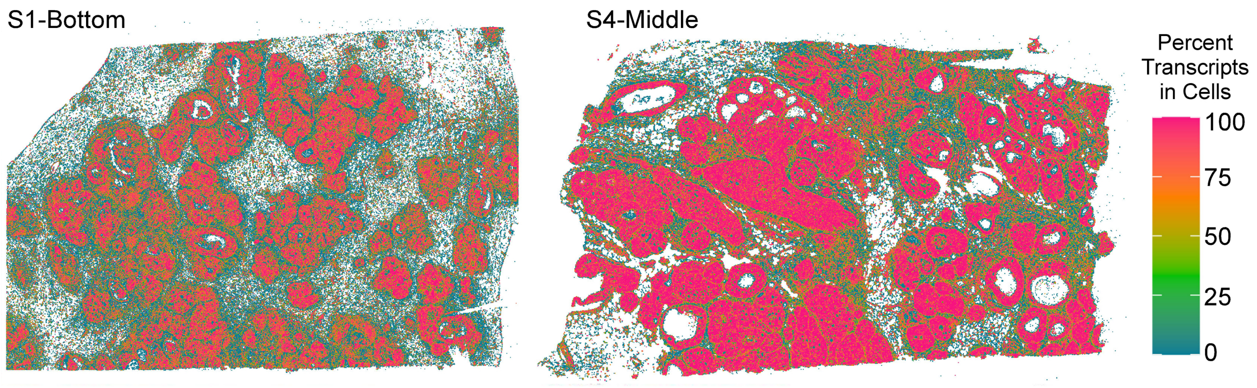


**Supplemental Figure S4. Cell segmentation is less reliable in the tumor periphery.** Spatial plots for two DCIS sections showing the percent transcripts assigned to cells using multimodal segmentation. The transcripts were partitioned into 10 x10 μm$^2$ bins. Bins with a transcript density <0.1/μm$^2$ were excluded.

### Supplemental Figure S5

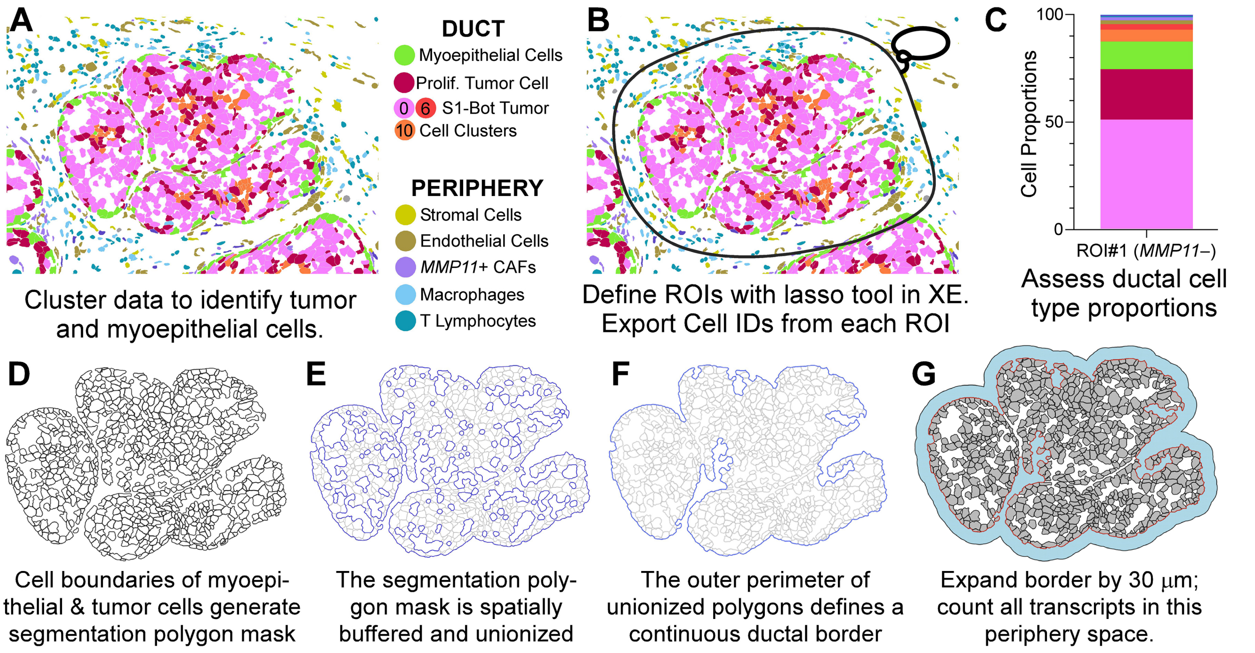


**Supplemental Figure S5. Strategy for defining the tumor periphery.** (**A**) Example duct and surrounding microenvironment from section S1-Bot showing annotated cell clusters including five distinct tumor subtypes (0, 2, 6, 8, 10) and a proliferative tumor cell cluster. (**B**) Manual lasso selection in Xenium Explorer (XE) was used to roughly outline each duct and peripheral cells, thus reducing the search area for downstream analysis. (**C, D**) Exported tumor and myoepithelial cell IDs were used to visualize the duct composition of each ROI and generate (**E**) a segmentation polygon mask. (**F**) Polygons are spatially buffered and unionized (merged together) to minimize gaps between adjacent cells and produce a continuous duct border. (**G**) The tumor periphery is defined by expanding the ductal border outward by 30 μm. In this example, no neighboring ducts overlap the periphery; in cases of overlap, those neighboring regions are excluded, with minor changes reflected in cell type proportion plots like **C**.

### Supplemental Figure S6

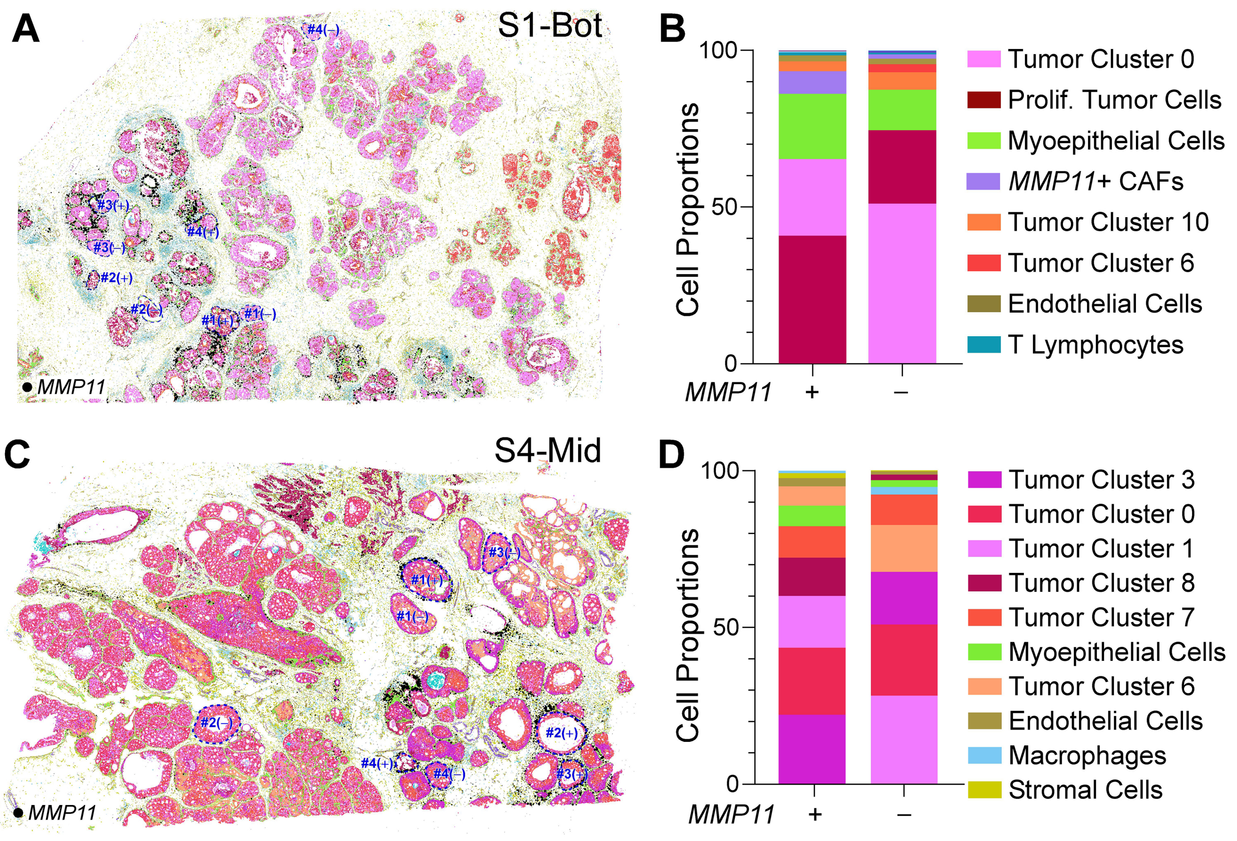


**Supplemental Figure S6. Selected ROIs for tumor periphery analysis and duct cell type composition.** (**A, B**) Spatial plots colored by cell type showing areas (blue dotted lines) selected for analysis of tumor periphery and duct composition. Each section has eight ROIs (four *MMP11*+ and four *MMP11*−) selected for analysis. *MMP11* quantitation for each of these ROIs is shown in Figure 2F. (**C, D**) Cell type composition of *MMP11*+ versus *MMP11*− ducts.
